# Activating FGFR1 restores Integrin-β1–mediated fibronectin sensing in satellite cells of aged mice

**DOI:** 10.64898/2026.02.18.706475

**Authors:** Tze-Ling Chang, Tenaya K. Vallery, Thea S. Zlatkov, Alicia A. Cutler, Jesse V. Kurland, Carson H. Butcher, Kristi S. Anseth, Bradley B. Olwin

**Author notes:** Correspondence (K.S.A.); (B.B.O.).

## Abstract

Muscle satellite cells (SCs), essential for skeletal muscle regeneration, decline in number and function with age, contributing to sarcopenia. A fully defined viscoelastic hydrogel that preserves SC-myofiber interactions and supports tunable densities of fibronectin-derived RGD ligands was used to investigate age-related defects in extracellular matrix sensing by SCs. Elevating RGD density increased the number of activating and proliferating SCs on myofibers from young mice, whereas SCs from aged mice were unresponsive. Loss of FGF receptor 1 signaling in SCs from aged mice abrogated the coordinated Syndecan-4 and Integrin-β1 matrix response observed in SCs from young mice. Activating Integrin-β1 promoted asymmetric division and self-renewal in SCs from young mice whereas combined FGFR1 and Integrin-β1 signaling drove symmetric expansion. In SCs from aged mice, FGFR1 dysfunction disrupted this balance, impairing asymmetric division, but constitutive FGFR1 activation restored receptor co-localization, self-renewal, and fibronectin responsiveness. Therefore, FGFR1 integrates matrix and growth factor signals, suggesting that targeting the FGFR1–Integrin-β1 axis may enhance SC regenerative potential in aging organisms.

## Introduction

Skeletal muscle satellite cells or stem cells (SCs), located between the basal lamina and the sarcolemma of muscle myofibers, are essential for regenerating muscle^1^. Upon injury, quiescent SCs activate and either asymmetrically or symmetrically divide to renew the stem cell pool or proliferate, respectively. Proliferating SCs or myoblasts express MyoD1 and can terminally differentiate and fuse to repair damaged myofibers^2^. During aging, SC numbers are reduced, proliferative capacity is diminished, and signaling responses are compromised, culminating in delayed or incomplete muscle repair^3–5^.

Age-associated declines in the responsiveness of fibroblast growth factor receptor 1 (FGFR1) compromise SC self-renewal, thereby impairing muscle maintenance and regeneration^6, 7^ (Cutler et al., unpublished). FGFR1 integrates extracellular cues with intracellular signaling pathways to prevent SC terminal differentiation and to promote SC asymmetric division^8^. In SCs from aged mice, a cell-autonomous loss of self-renewal occurs as FGFR1 ligand responsiveness is reduced and compensatory p38 MAPK signaling is elevated^6, 9^. Constitutively activating FGFR1 in cultured SCs restores the self-renewal and engraftment capacity of SCs from aged mice^6^, while constitutively activating an SC-specific inducible FGFR1 (caFGFR1) *in vivo* rescues cell-intrinsic aging-related defects and promotes asymmetric division in SCs from aged mice (Cutler et al., unpublished).

FGFR1 signaling in SCs requires Syndecan-4 and Integrin-β1 as ablating either gene abrogates FGFR1 signaling in the absence of Syndecan-4^10^ or reduces FGFR1 responsiveness to FGFs in the absence of Integrin-β1^11^. Syndecan-4, a transmembrane heparan sulfate proteoglycan, is required for FGFR1 to bind FGF and activate the receptor^12–14^. Integrin-β1, a cell surface transmembrane receptor, is essential for SC responsiveness to FGF-2. Deletion of either Syndecan-4 or Integrin-β1 alters FGF signaling, reducing SC expansion upon injury and eliminating SC self-renewal^10, 11^. In SCs from aged mice, Integrin-β1 signaling is impaired, and restoring Integrin-β1 activity rescues FGF responsiveness and improves regeneration^11^; however, how coordinated signaling from Syndecan-4, Integrin-β1 and FGFR1 are compromised in SCs as mice age is poorly understood.

The ECM surrounding SCs undergoes dynamic remodeling^15^ to support fibronectin-driven symmetric SC expansion^16, 17^. Fibronectin, which binds integrins, accumulates around SCs following injury, promoting Wnt7a signaling via Syndecan-4 binding^18^. In the absence of fibronectin, SC responsiveness to FGF-2 is reduced^11^. Thus, during aging, the ability of SCs to sense and respond to fibronectin is compromised, affecting FGF signaling and impairing SC self-renewal. We developed and employed a fully defined synthetic hydrogel matrix to better understand interactions between FGFR1, Integrin-β1, and Syndecan-4, and how aging influences these interactions. A material platform with tailorable density of fibronectin-derived epitope (RGD) provides muscle-relevant viscoelastic microenvironments for myofiber culture^19^. The hydrogel platform maintains SC polarity and preserves undifferentiated SCs on encapsulated myofibers while providing a SC-myofiber matrix interface that prevents myofiber hypercontraction.

SCs on myofibers from aged mice cultured in the hydrogel matrix respond poorly to RGD and fail to polarize FGFR1. In contrast, ectopically activating Integrin-β1 in SCs from young mice promotes SC self-renewal, while combined ectopic activation of Integrin-β1 and ectopic activation of FGFR1 stimulates SC symmetric expansion. In SCs from aged mice, impaired FGFR1 signaling abrogates sensitivity of the cells to RGD. Constitutively activating FGFR1 in SCs from aged mice restores FGFR1 polarization, co-localization of Integrin-β1 and Syndecan-4 with FGFR1, as well as RGD ligand sensitivity that cumulatively promote asymmetric division. At high RGD density in the hydrogel matrix, ectopically activating FGFR1 promotes SC symmetric expansion, increasing myoblast proliferation that supports muscle regeneration.

## Results

### Aging diminishes responsiveness of SCs on myofibers to RGD density in hydrogel environments

Viscoelastic hydrogel formulation used for myofiber encapsulation prevents myofiber hypercontraction, maintains SC motility, and preserves non-differentiated SCs^19^. Moreover, encapsulating myofibers in hydrogels with high RGD density promotes SC activation and proliferation. How SCs on myofibers from aged mice respond to hydrogels with varying RGD density remains unclear. Since viscoelastic hydrogels containing 88% alkyl-hydrazone crosslinks maintain SC quiescence in myofiber culture^19^, we used the same formulation for this study. The hydrogel chemistry and mechanical properties are shown in **Figure S1** and **Table S1**.

Myofibers isolated from young (4-6 months old) and aged (20-24 months old) mice were encapsulated in hydrogels containing 0.1 mM, 0.5 mM, or 1 mM RGD **(Figure 1A)**. Encapsulated myofibers were cultured for two days, with a 2-hour EdU pulse administered prior to fixation to quantify proliferating SCs. SC fate was assessed by Pax7 and MyoD immunoreactivity **(Figures 1B and 1C)**. On myofibers from young mice, the percentage of Pax7⁺ cells increased from 50 ± 3% in hydrogels with 0.1 mM RGD to 66 ± 3% and 61 ± 3% in hydrogels with 0.5 mM and 1 mM RGD, respectively **(Figure 1D)**. In contrast, fewer than 35% of cells on myofibers from aged mice were Pax7^+^, regardless of RGD concentration. Among Pax7^+^ cells from young mice, 52 ± 2% were MyoD^+^ when cultured in hydrogels with 0.1 mM RGD, increasing to over 70% in 0.5 mM and 1 mM RGD **(Figure 1E)**. However, on myofibers from aged mice, fewer than 35% of Pax7^+^ cells were MyoD^+^ regardless of RGD concentration. In addition, 22 ± 4% of Pax7^+^ cells on myofibers from young mice were EdU^+^ in 0.1 mM RGD hydrogels, increasing to 41 ± 4% and 58 ± 4% in 0.5 mM and 1 mM RGD, respectively **(Figure 1F)**. In contrast, fewer than 30% of Pax7^+^ cells from aged mice were EdU^+^, with no increase in response to RGD concentration. Compared to myofibers from young mice, myofibers from aged mice exhibited significantly fewer Pax7^+^ cells, indicating reduced SC proliferation. While SC activation and proliferation on myofibers from young mice increased with RGD density, SCs from aged mice were unresponsive, suggesting impaired sensitivity to environmental cues.

**Figure 1.**
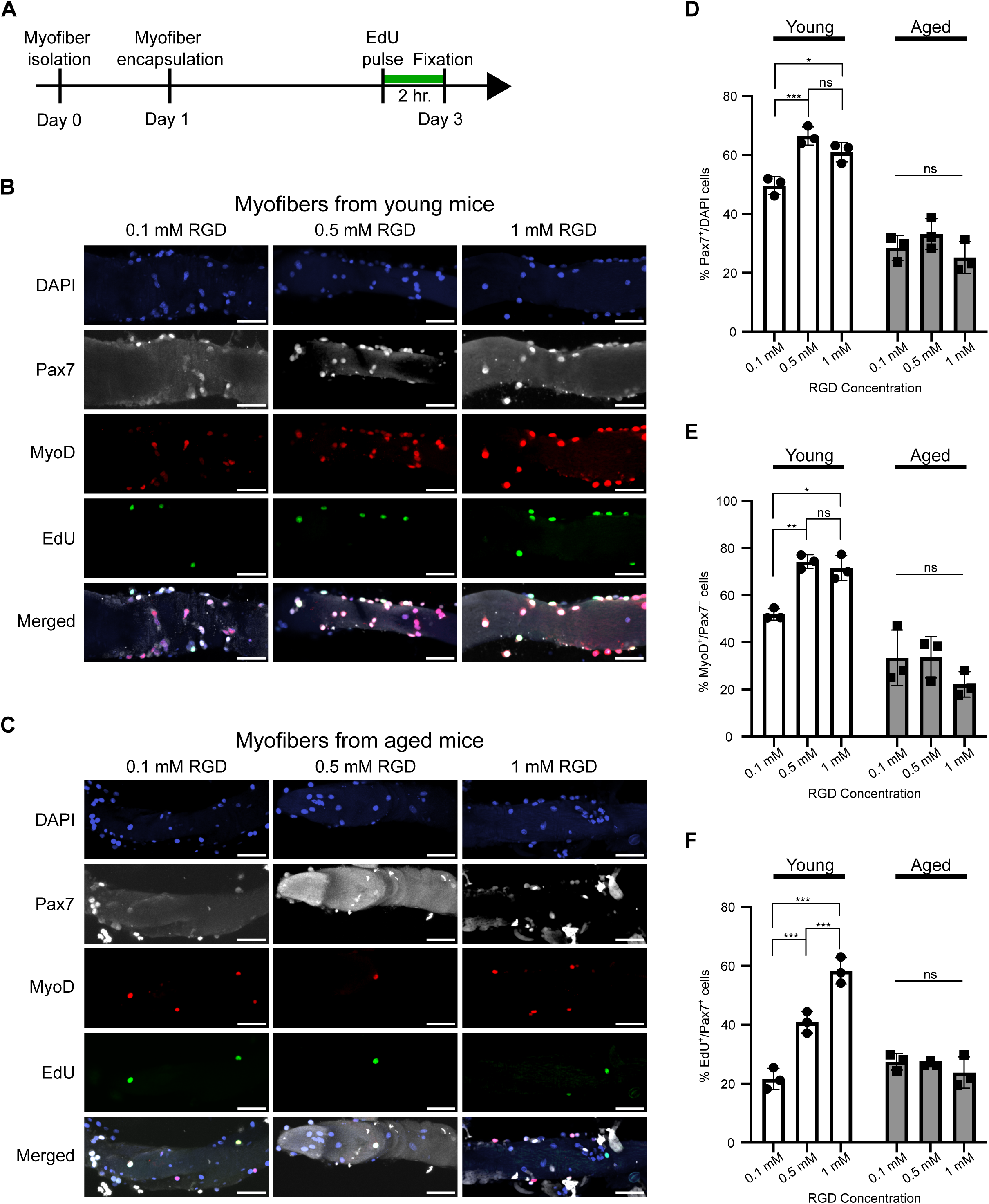
SCs on myofibers from aged mice lose sensitivity to RGD density in hydrogels. (A) Experimental scheme of encapsulating myofibers isolated from young (4-6 months old) and aged mice (20-24 months old) in hydrogels. (B and C) Representative images of myofibers from young (B) and aged (C) mice embedded in hydrogels with 0.1 mM, 0.5 mM, and 1 mM RGD and labeled with Pax7, EdU, and MyoD after 2 days of culture. Scale bars, 50 μm. (D-F) Quantification of Pax7^+^ (D), MyoD^+^ (E), and EdU^+^ (F) for SCs on myofibers embedded in hydrogels with 0.1 mM, 0.5 mM, and 1 mM RGD. *n* > 9 myofibers analyzed with samples from 3 mice, and statistics performed based on 3 independent mice; data are presented as mean ± SD; *\*P* < 0.05, *\*\*P* < 0.01, and *\*\*\*P* < 0.001 in a two-way ANOVA test.

### Syndecan-4 and integrins are predicted key receptors mediating fibronectin signaling in SCs based on single-cell and single-nucleus RNA sequencing

To identify key receptors involved in fibronectin signaling in SCs, single-cell and single-nucleus RNA sequencing data from young and aged mice, both uninjured and 4 days post-injury, were processed and clustered by cell type with *CellRanger* and *Seurat*. Cell signaling interactions were inferred using *CellChat. CellChat* analysis of fibronectin signaling predicts *Sdc4* as the major receptor of fibronectin expressed by SCs from young and aged uninjured mice **(Figure 2A)**. Upon injury, myogenic progenitor cells and SCs in both young and aged mice, express *Cd44* and fibronectin-interacting integrins, including *Itgav*, *Itgb6*, and *Itgb1*.

**Figure 2.**
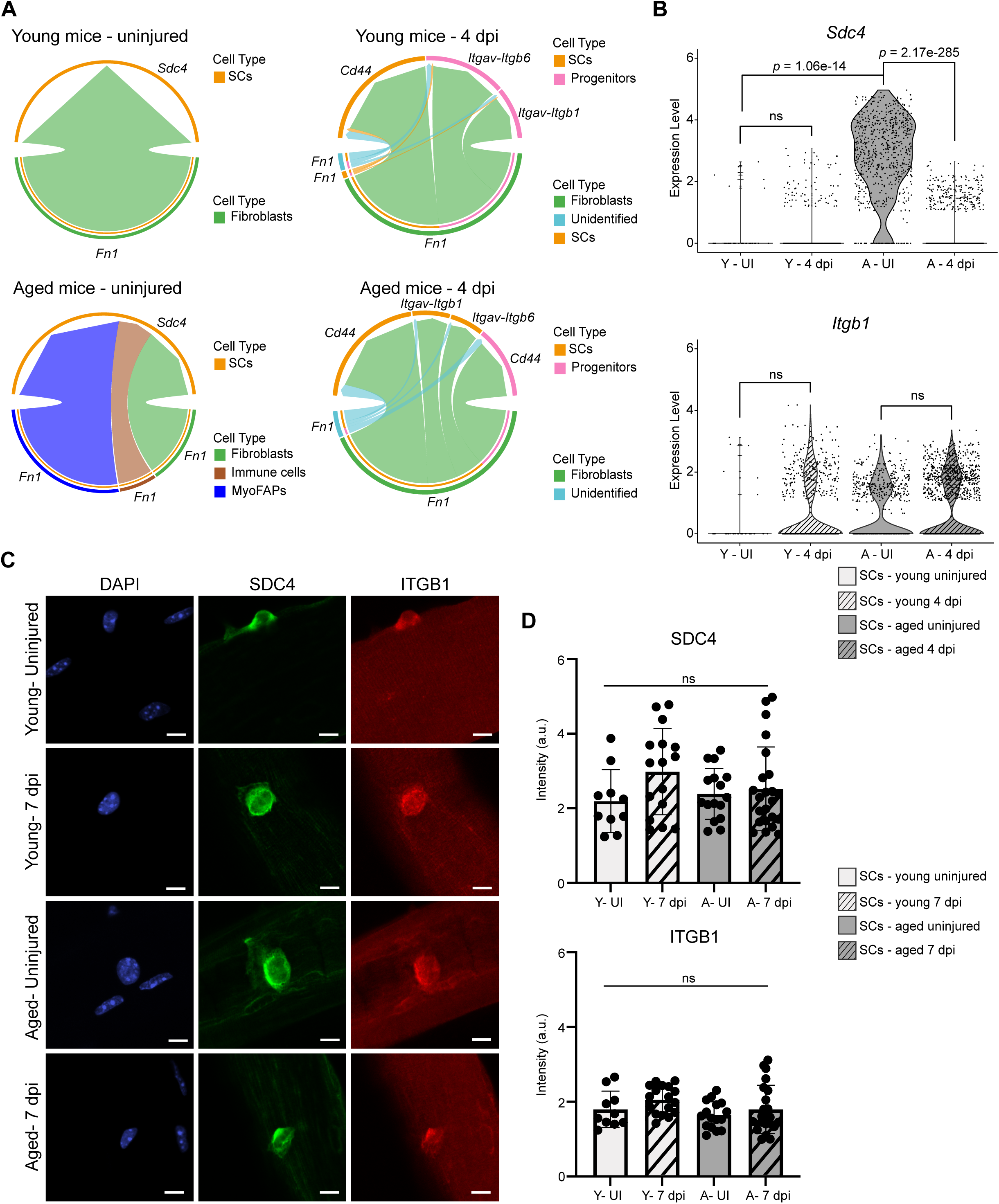
Syndecan-4 and Integrin-β1 mediate the interaction between SCs and fibronectin within the SC niche in both young and aged mice. (A) *CellChat* chord plots of the inferred fibronectin signaling network from single-cell and single-nucleus RNA sequencing data collected from young and aged mouse muscle, both uninjured and at 4 days post-injury (dpi). Ligands expressed by sender cell type groups are listed on the bottom while receptors expressed by receiver cell type groups are displayed on the top. (B) Violin plots of transcript levels of *Sdc4* and *Itgb1* in SCs from single-cell and single-nucleus sequencing data from young and aged mice, both uninjured and at 4 dpi. Statistical significance was measured by *Seurat’s* Wilcox rank sum test and is represented by p. (C) Representative images of SCs on myofibers from EDL muscles of uninjured young and aged mice, and at 7 dpi, immunoreactive for Syndecan-4 and Integrin-β1. Scale bars, 10 μm. (D) Quantification of protein levels for Syndecan-4 (SDC4) and Integrin-β1 (ITGB1) in SCs on uninjured myofibers from young and aged mice, and at 7 dpi. *n* > 10 myofibers analyzed with samples from 3 mice; data are presented as mean ± SD; statistics performed in a one-way ANOVA test.

To validate *CellChat* predictions, we examined expression levels of *Sdc4* and *Itgb1* in single-cell and single-nucleus RNA sequencing (sc/snRNA-seq) data. *Sdc4* is expressed at low levels in SCs from young mice, both in uninjured SCs and in SCs following an injury **(Figure 2B)**. In contrast, *Sdc4* is significantly upregulated in SCs from aged uninjured mice, but its expression drops markedly following injury. Fewer SCs from young uninjured mice express *Itgb1* compared SCs from uninjured aged mice or injured young or aged mice; however, the differences in expression levels among all experimental conditions are not statistically significant.

In addition to transcript levels, we examined protein levels of Syndecan-4 and Integrin-β1 in SCs on myofibers isolated from EDL muscles of uninjured young and aged mice, and from EDL muscles of injured young and aged mice at 7 days post-injury **(Figure 2C)**. Protein expression was quantified by measuring the mean fluorescence intensity on the cell membrane, subtracting the background signal defined as the mean intensity within the myofiber, and then normalizing the result to the mean nuclear DAPI intensity **(Figure 2D)**. Surprisingly, we observed no significant differences in the cell surface protein levels of Syndecan-4 and Integrin-β1 across the four experimental conditions, suggesting that upregulation of their transcript levels in aged uninjured mice may reflect a compensatory response to impaired downstream signaling. Given that Syndecan-4 and Integrin-β1 mediate the fibronectin response in SCs, we next investigated additional receptors that associate with Syndecan-4 and Integrin-β1.

### Constitutive activation of FGFR1 partially rescues the dysregulation of SC activation in aged mice

Given the lack of significant differences in protein levels for Syndecan-4 and Integrin-β1, we next investigated whether changes in FGFR1 might contribute to the impaired responsiveness to RGD signals. FGFR1 signaling is attenuated in SCs from aged mice^6^ and deletion of either Syndecan-4 or Integrin-β1 compromises the SC response to FGF stimulation^10, 11^. Since Syndecan-4 is a co-receptor for FGFR1^20, 21^, Integrin-β1 may form a ternary complex with FGFR1 and Syndecan-4 to mediate environmental signal transduction. We assessed FGFR1 protein levels by FGFR1 immunoreactivity in SCs on myofibers isolated from EDL muscles of uninjured young and aged mice, and 7 days post-injury **(Figure 3A)**. FGFR1 protein was upregulated in SCs from young mice 7 days post-injury compared to uninjured controls **(Figure 3B)**. In contrast, in SCs from aged mice, we observed lower FGFR1 expression, with no observable upregulation following injury, suggesting dysregulated FGFR1 expression in SCs from aged mice.

**Figure 3.**
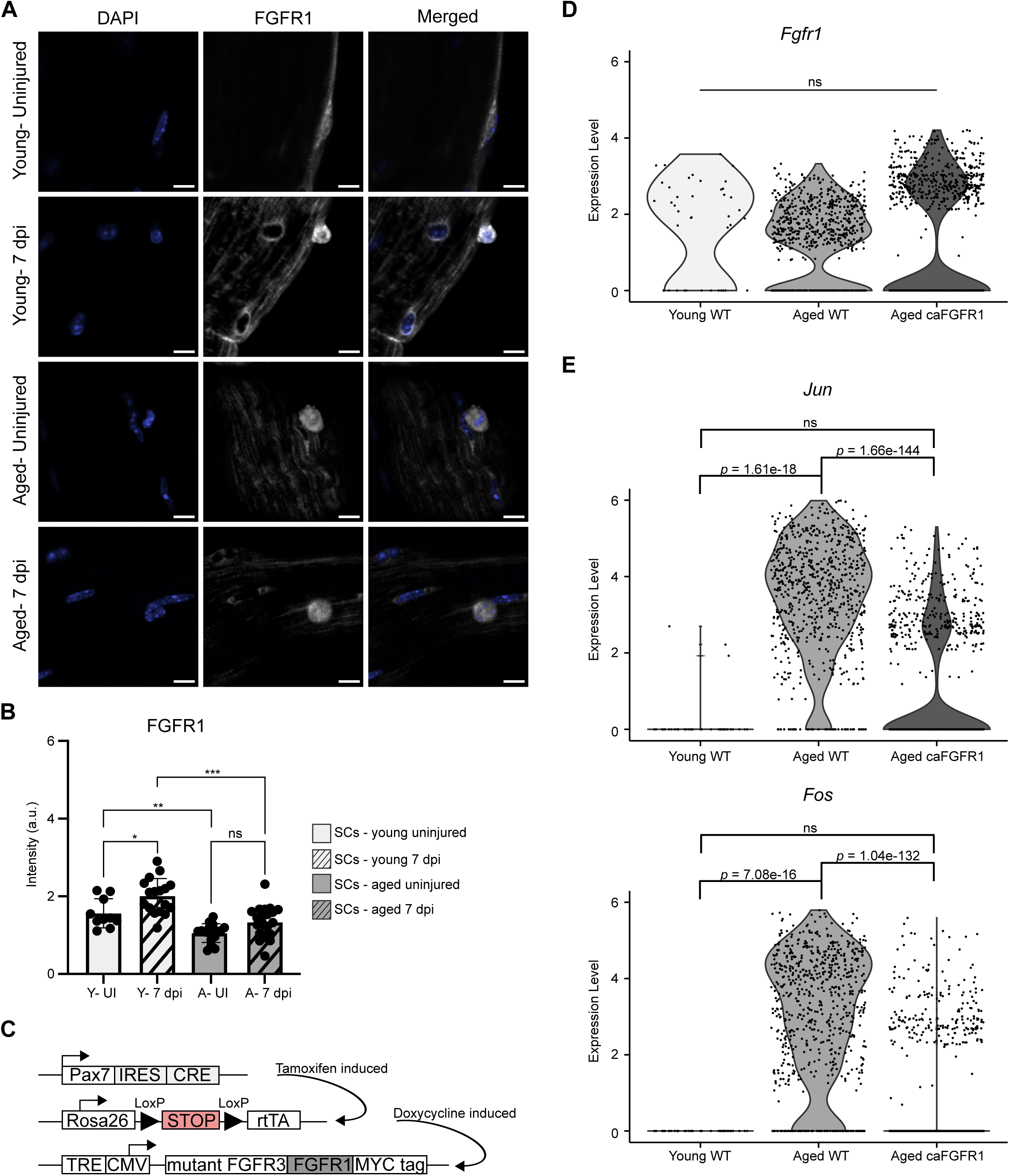
Constitutively active FGFR1 partially rescues the dysregulation of SC activation in aged mice. (A) Representative images of SCs on myofibers from young and aged mice, both uninjured and at 7 dpi, assessed for FGFR1 immunoreactivity. Scale bars, 10 μm. (B) Quantification of protein levels of FGFR1 for SCs on uninjured myofibers from young and aged mice, and at 7 dpi. *n* > 10 myofibers analyzed with samples from 3 mice; data are presented as mean ± SD; *\*P* < 0.05, *\*\*P* < 0.01, and *\*\*\*P* < 0.001 in a one-way ANOVA test. (C) Schematic diagram of caFGFR1 mouse genetics. (D) Violin plot of transcript levels of *Fgfr1* from single-cell and single-nucleus RNA sequencing data of SCs from young wild type, aged wild type, and aged caFGFR1+ mice. Statistical significance was measured by *Seurat’s* Wilcox rank sum test. (E) Violin plots of *Jun* and *Fos* transcript levels in SCs from young wild type, aged wild type, and aged caFGFR1+ mice. Statistical significance was measured by *Seurat’s* Wilcox rank sum test and is represented by p.

To rescue the impaired FGFR1 function in SCs from aged mice, mice were bred for temporally inducible, constitutively active FGFR1 (caFGFR1) expression specifically in SCs via the tetracycline-responsive element (Cutler et al., unpublished) **(Figure 3C)**. Briefly, the caFGFR1 transgene encodes a chimeric receptor combining the FGFR3c(R248C) extracellular and transmembrane domains with the FGFR1 intracellular kinase domain and a C-terminal MYC tag and is expressed only in the presence of doxycycline^22^ (Cutler et al., unpublished). To further validate FGFR1 expression in the caFGFR1+ mice, we examined transcript levels of *Fgfr1* in single-cell and single-nuclear RNA sequencing data, and no statistically significant differences in *Fgfr1* transcript levels were observed among the three groups **(Figure 3D)**, indicating that the ectopically expressed caFGFR1 is expressed at levels similar to wild type. Incomplete recombination efficiency as well as feedback control on FGFR1 expression may account for the lack of transgene overexpression (Cutler et al., unpublished). We then tested whether activating FGFR1 expression in SCs from aged mice influences the gene expression associated with the fibronectin signaling pathway. The downstream targets *Jun* and *Fos*, which are typically upregulated in activated SCs, are significantly elevated in SCs from aged wild-type mice compared to those from young wild-type mice **(Figure 3E)**. Surprisingly, ectopically activating FGFR1 in SCs from aged mice reduces *Jun* and *Fos* transcript levels, restoring their expression to levels comparable to those in young mice **(Figure 3E)**.

### Constitutive activation of FGFR1 restores FGFR1 polarization and co-localization of Syndecan-4 and Integrin-β1 with FGFR1 in SCs from aged mice

Upon activation, FGFR1 polarizes at the SC membrane and asymmetrically activates p38 mitogen-activated protein kinase (MAPK), regulating SC self-renewal^6^. To investigate if caFGFR1 in SCs from aged mice rescues asymmetric FGFR1 distribution, myofibers were isolated from young caFGFR1–, aged caFGFR1–, and aged caFGFR1+ mice (**Figure 4A**). All mice received daily tamoxifen injections for two consecutive days, followed by three days of doxycycline chow prior to myofiber isolation. Isolated myofibers were cultured in suspension with FGF-2 for 36 hours, fixed, and then assessed for Syndecan-4 immunoreactivity to identify SCs, for endogenous FGFR1 immunoreactivity with an anti-FGFR1 extracellular domain antibody, and for caFGFR1 immunoreactivity using an anti-FGFR3 extracellular domain antibody (**Figure 4B**). Since *Fgfr3* is not expressed in SCs (**Figures S2A and S2B**) and recombined SCs from caFGFR1+ mice express an FGFR3 extracellular domain fused to the FGFR1 intracellular domain^22^, FGFR3 immunoreactivity identifies caFGFR1 protein. A significantly lower proportion of Syndecan-4^+^ SCs from aged caFGFR1– mice polarized FGFR1 compared with those from young caFGFR1– mice (**Figure 4C**), in agreement with prior published data^6^. In contrast, ectopically expressing caFGFR1 in SCs from aged mice restored polarized FGFR1 distribution that appeared similar to FGFR1 polarization in SCs from young mice.

**Figure 4.**
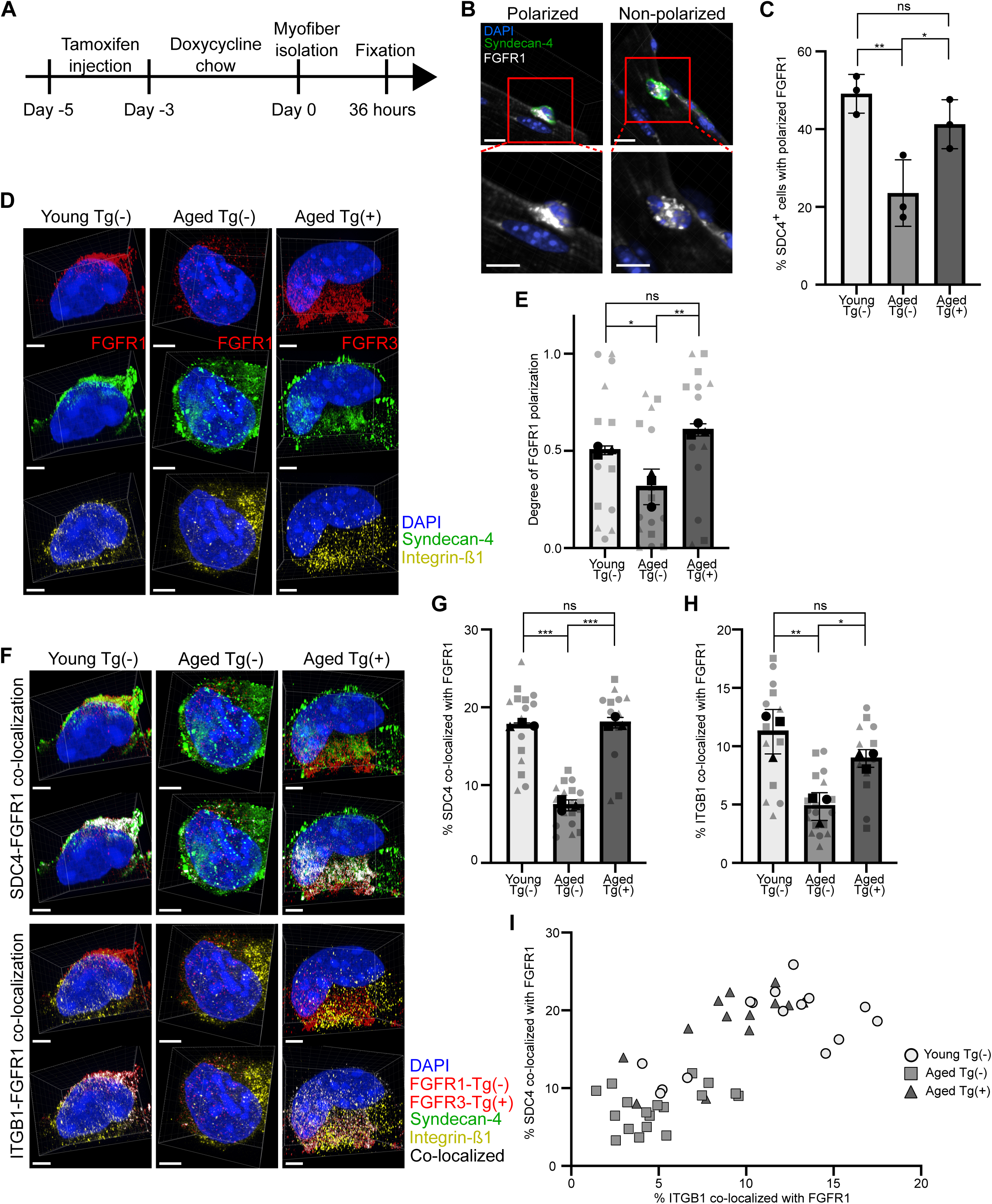
Constitutively active FGFR1 restores FGFR1 polarization and Syndecan-4 and Integrin-β1 co-localization with FGFR1 in SCs from aged mice. (A) Experimental scheme showing treatment of young caFGFR1–, aged caFGFR1–, and aged caFGFR1+ mice with tamoxifen and doxycycline chow, followed by myofiber isolation and culture with fibroblast growth factor-2 (FGF-2) for 36 hours prior to fixation. (B) Representative images of polarized and non-polarized FGFR1 in SCs on myofibers. Scale bars, 10 μm. (C) Percentage of Syndecan-4^+^ cells with polarized FGFR1 on myofibers from young caFGFR1–, aged caFGFR1–, and aged caFGFR1+ mice. *n* > 60 Syndecan-4^+^ cells analyzed with samples from 3 mice, and statistics performed based on 3 independent mice; data are presented as mean ± SD *\*P* < 0.05 and *\*\*P* < 0.01 in a one-way ANOVA test. (D) Representative photo-expansion microscopy images of SCs from young caFGFR1–, aged caFGFR1–, and aged caFGFR1+ mice, immunoreactive for Syndecan-4, Integrin-β1, and either the endogenous FGFR1 extracellular domain in unrecombined SCs or the mutant FGFR3 extracellular domain in recombined SCs. Scale bars, 2 μm. (E) Degree of FGFR1 polarization in Syndecan-4^+^ SCs from young caFGFR1–, aged caFGFR1–, and aged caFGFR1+ mice. *n* > 10 Syndecan-4^+^ cells analyzed (represented by shaded data points) with samples from 3 mice (solid symbols denoted by circles, squares, and triangles), and statistics performed based on 3 independent mice; data are presented as mean ± SD; *\*P* < 0.05 and *\*\*P* < 0.01 in a one-way ANOVA test. (F) Representative images of FGFR1 co-localized with Syndecan-4 and Integrin-β1 in SCs from young caFGFR1–, aged caFGFR1–, and aged caFGFR1+ mice. Scale bars, 2 μm. (G and H) Percentage of Syndecan-4 (G) and Integrin-β1 (H) co-localized with FGFR1 in SCs from young caFGFR1–, aged caFGFR1–, and aged caFGFR1+ mice. *n* > 10 Syndecan-4 ^+^ cells analyzed (represented by shaded data points) with samples from 3 mice (solid symbols denoted by circles, squares, and triangles), and statistics performed based on 3 independent mice; data are presented as mean ± SD; *\*P* < 0.05, *\*\*P* < 0.01, and *\*\*\*P* < 0.001 in a one-way ANOVA test. (I) Correlation of Syndecan-4-FGFR1 and Integrin-β1-FGFR1 colocalization in SCs from young caFGFR1–, aged caFGFR1–, and aged caFGFR1+ mice.

FGFR1 activation requires coordinated interactions with multiple membrane receptors, including Syndecan-4^10, 20, 21^ and Integrin-β1^12^. However, visualization of membrane receptor organization on SCs is technically challenging due to their small size and the limited resolution of conventional confocal microscopy. To assess FGFR1 polarization and receptor co-localization at high spatial resolution, photo-expansion microscopy (PhotoExM)^23^ was utilized to visualize membrane receptor distribution in three dimensions. SCs on isolated myofibers were fixed and assessed for immunoreactivity without permeabilization to selectively label extracellular domains. SCs from young and aged caFGFR1– mice were assessed for cell surface immunoreactivity of Syndecan-4, Integrin-β1, and the endogenous FGFR1 extracellular domain, whereas SCs from aged caFGFR1+ mice were assessed for immunoreactivity of Syndecan-4, Integrin-β1, and the FGFR3 extracellular domain. Immunolabeled myofibers were then embedded in polyelectrolyte expansion hydrogels and subsequently expanded in water to achieve a 4.5X linear expansion, enabling higher-resolution imaging (**Figure 4D**).

To determine the degree of FGFR1 polarization, a plane passing through the cell’s center of mass and perpendicular to the myofiber axis was defined (**Figure S3A**). The volume of FGFR1 on either side of the defined plane was quantified and expressed as a ratio of the total FGFR1 volume within the cell. This ratio was then normalized to a scale from 0 to 1 to define the degree of polarized FGFR1. The degree of polarized FGFR1 decreased from 0.50 ± 0.02 in SCs from young caFGFR1– mice to 0.32 ± 0.09 in those from aged caFGFR1– mice; however, polarized FGFR1 increased to 0.61 ± 0.03 in SCs from aged caFGFR1+ mice, further indicating that constitutive activation of FGFR1 restores their polarization ability (**Figure 4E**).

Next, we analyzed Syndecan-4 and Integrin-β1 co-localization with FGFR1 using the Imaris Analysis Workstation. SCs from young and aged caFGFR1– mice were assessed for endogenous FGFR1 extracellular domain using anti-FGFR1 antibodies, and recombined SCs from aged caFGFR1+ mice were assayed for caFGFR1 with antibodies to the extracellular FGFR3 domain. The red, green, and yellow channels represent FGFR1/FGFR3, Syndecan-4, and Integrin-β1, respectively, while the white channel indicates regions where either Syndecan-4 or Integrin-β1 co-localizes with FGFR1/FGFR3 (**Figure 4F**). In SCs from young caFGFR1– mice, 17.7 ± 0.3% of Syndecan-4 co-localized with FGFR1, while only 7.4 ± 0.7% of Syndecan-4 co-localized with FGFR1 in SCs from aged caFGFR1– mice (**Figure 4G**). In SCs from aged caFGFR1+ mice, 18.0 ± 0.7% of Syndecan-4 colocalized with caFGFR1, restoring co-localization to that observed for FGFR1 in young mice (**Figure 4G**). A similar pattern was observed with Integrin-β1, where co-localization with FGFR1 decreased from 11.3 ± 1.9% in SCs from young caFGFR1– mice to 4.8 ± 1.2% in SCs from aged caFGFR1– mice, and was subsequently restored to 9.0 ± 0.9% in SCs from aged caFGFR1+ mice (**Figure 4H**). The scatter plot presents the correlation between Syndecan-4 (y-axis) or Integrin-β1 (x-axis) co-localization with FGFR1, with each point representing a single SC measurement from the indicated groups (**Figure 4I** and **Figure S3B**). SCs from young caFGFR1– and aged caFGFR1+ mice exhibited a relatively high percentage of Integrin-β1 and Syndecan-4 puncta co-localized with FGFR1, clustering predominantly in the upper-right quadrant of the plot. In contrast, SCs from aged caFGFR1– mice showed reduced co-localization, with values clustered in the lower-left quadrant. These results suggest that activating FGFR1 restores both Integrin-β1 and Syndecan-4 co-localization with FGFR1 in SCs from aged mice.

### Integrin-β1 modulates asymmetric division in an FGF-dependent manner in SCs on myofibers from young mice

Based on the observation that Integrin-β1 co-localizes with FGFR1 when FGFR1 is activated, we next asked if Integrin-β1 and FGFR1 cooperate in regulating SC behavior. Young caFGFR1– mice received daily tamoxifen injections for two consecutive days, followed by three days of doxycycline chow prior to myofiber isolation (**Figure 5A**). Isolated myofibers were then cultured in suspension with or without FGF-2, in the presence of Integrin-β1-blocking antibody AIIB2 or Integrin-β1-activating antibody TS2/16, for 36 hours before fixation. SCs on myofibers were assayed for Syndecan-4 immunoreactivity to identify SCs (**Figure 5B**), and the number of Syndecan-4^+^ cells on myofibers was quantified (**Figure 5C**). With FGF stimulation, 8.7 ± 0.4 Syndecan-4^+^ cells per millimeter of myofiber were present, decreasing to 3.4 ± 0.4 with AIIB2 treatment and increasing to 17.9 ± 3.9 with TS2/16 treatment. In contrast, in the absence of FGF, only 4.1 ± 2.5 Syndecan-4^+^ cells per millimeter of myofiber were present, whether or not Integrin-β1 was blocked with AIIB2 treatment. Activating Integrin-β1 with TS2/16 markedly increased SC numbers per millimeter of myofiber to 13.1 ± 2.3. Thus, activating Integrin-β1 in the absence of FGF is sufficient to maintain SC numbers; however, simultaneously activating FGFR1 and Integrin-β1 dramatically expands the SC population.

**Figure 5.**
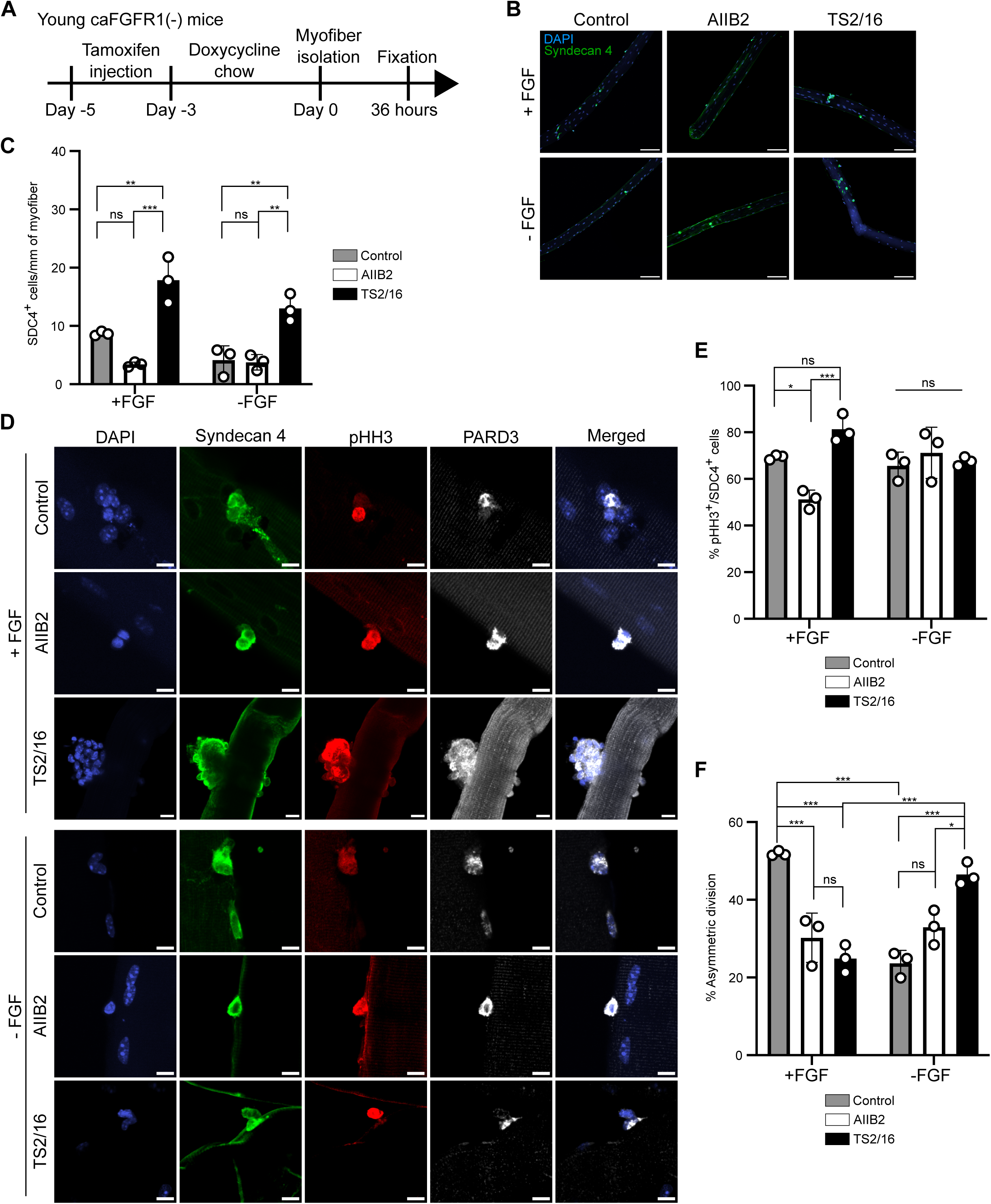
Integrin-β1 modulates FGFR1-dependent asymmetric division in SCs on myofibers from young mice. (A) Experimental scheme for treatment of young caFGFR1– mice with tamoxifen and doxycycline chow, followed by myofiber isolation and culture prior to fixation. (B) Representative images of myofibers cultured with or without FGF-2, in the presence of AIIB2 or TS2/16, where SCs were identified by Syndecan-4 immunoreactivity. Scale bars, 100 μm. (C) Number of Syndecan-4^+^ cells on myofibers cultured with or without FGF-2, in the presence of AIIB2 or TS2/16. *n* > 10 myofibers analyzed with samples from 3 mice, and statistics performed based on 3 independent mice; data are presented as mean ± SD; *\*\*P* < 0.01 and *\*\*\*P* < 0.001 in a two-way ANOVA test. (D) Representative images of SCs on myofibers cultured with or without FGF-2, in the presence of AIIB2 or TS2/16, identified by Syndecan-4 immunoreactivity and assessed for phospho-histone H3 (pHH3) and PARD3 immunoreactivity. Scale bars, 10 μm. (E and F) Percentage of pHH3^+^ immunoreactive cells (E) and percentage of asymmetric division (F) on myofibers cultured with or without FGF-2, in the presence of AIIB2 or TS2/16. *n* > 200 Syndecan-4^+^ cells analyzed with samples from 3 mice, and statistics performed based on 3 independent mice; data are presented as mean ± SD; *\*P* < 0.05 and *\*\*\*P* < 0.001 in a two-way ANOVA test.

To further understand how FGFR1 and Integrin-β1 synergize to increase SC numbers, we examined division of individual Syndecan-4^+^ cells. Phospho-histone H3 (pHH3) immunoreactivity identifies cells undergoing mitosis (**Figure 5D**). In the presence of FGF, 69 ± 1% of Syndecan-4^+^ cells were pHH3^+^, decreasing to 51 ± 4% when Integrin-β1was inhibited with AIIB2 (**Figure 5E**). If Integrin-β1 was stimulated with TS2/16 in the presence of FGF, pHH3^+^ cells increased to 81 ± 6%, greater than with FGF alone, and pHH3^+^/Syndecan-4^+^ cells were seen clustering on the myofibers (**Figure 5D, E**). In the absence of FGF, no statistically significant differences were observed among all three groups (**Figure 5E**).

Ectopically activated FGFR1 polarizes and drives asymmetric division^6^, and thus, we asked if the FGFR1 polarity alters the extent of SC asymmetric division. During symmetric division, PARD3 is symmetrically distributed between the two daughter cells, whereas during asymmetric division, PARD3 is polarized in only one daughter cell that maintains the myogenic lineage, while the other daughter lacking PARD3 self-renews as a stem cell and re-acquires quiescence^8, 24^. In the presence of FGF, 52 ± 1% of SCs on intact myofibers divided asymmetrically, decreasing to 30 ± 6% when Integrin-β1 was inhibited by AIIB2 treatment (**Figure 5F**). Activating Integrin-β1 with TS2/16 surprisingly failed to restore asymmetric division when FGF was present. Given the high pHH3⁺ levels and presence of cell clusters, simultaneous activation of FGFR1 and Integrin-β1 appears to promote a shift toward symmetric division, resulting in an increase in cell division (**Figure S4A**). Without FGF, 24 ± 3% of cells divided asymmetrically, which was unaffected by AIIB2 treatment. Asymmetric division in the absence of FGF increased to 47 ± 3% with TS2/16 treatment. Since activating FGFR1 without stimulating integrin-ß1 dramatically increases asymmetric division^6^ (Cutler et al., unpublished) and activating Integrin-β1 in the absence of added FGF-2 increases asymmetric division, both FGFR1 and Integrin-β1 likely cooperate in regulating SC decisions to symmetrically divide or self-renew by asymmetrically dividing.

### Constitutive activation of FGFR1 partially rescues Integrin-β1-mediated regulation of asymmetric division in SCs from aged mice

Activating FGFR1 in aged mice polarizes Syndecan-4 and restores asymmetric division^6^ (Cutler et al., unpublished), and thus, we asked if activating FGFR1 would restore Integrin-β1 ability to regulate asymmetric division in SCs from aged mice as occurs in young mice. Aged caFGFR1– and caFGFR1+ mice received tamoxifen injections, followed by doxycycline chow prior to myofiber isolation (**Figure 6A**). Isolated myofibers were cultured in the presence of Integrin-β1-blocking antibody AIIB2 or Integrin-β1-activating antibody TS2/16 for 36 hours, and then fixed. SCs on myofibers were identified by Syndecan-4 immunoreactivity (**Figure 6B**) and quantified, where myofibers from aged caFGFR1– mice possessed 3.6 ± 0.7 SCs/mm of myofiber, with no significant change in SC numbers following AIIB2 treatment (**Figure 6C**). Treatment with TS2/16 increased SC number to 12.1 ± 3.2 per millimeter of myofiber, demonstrating that activating Integrin-β1 in myofibers from aged mice increases SC numbers. Myofibers from aged caFGFR1+ mice possessed 10.4 ± 0.7 SCs/mm of myofiber that decreased to 4.2 ± 1.0 SCs/mm of myofiber following AIIB2 treatment. A marked increase in Syndecan-4^+^ SCs to 24.7 ± 5.3 per millimeter of myofiber occurred following TS2/16 treatment, demonstrating potential synergy upon activating FGFR1 and Integrin-β1 for expanding SCs, similar to the data for young mice with added FGF-2 (**Figure 5C**).

**Figure 6.**
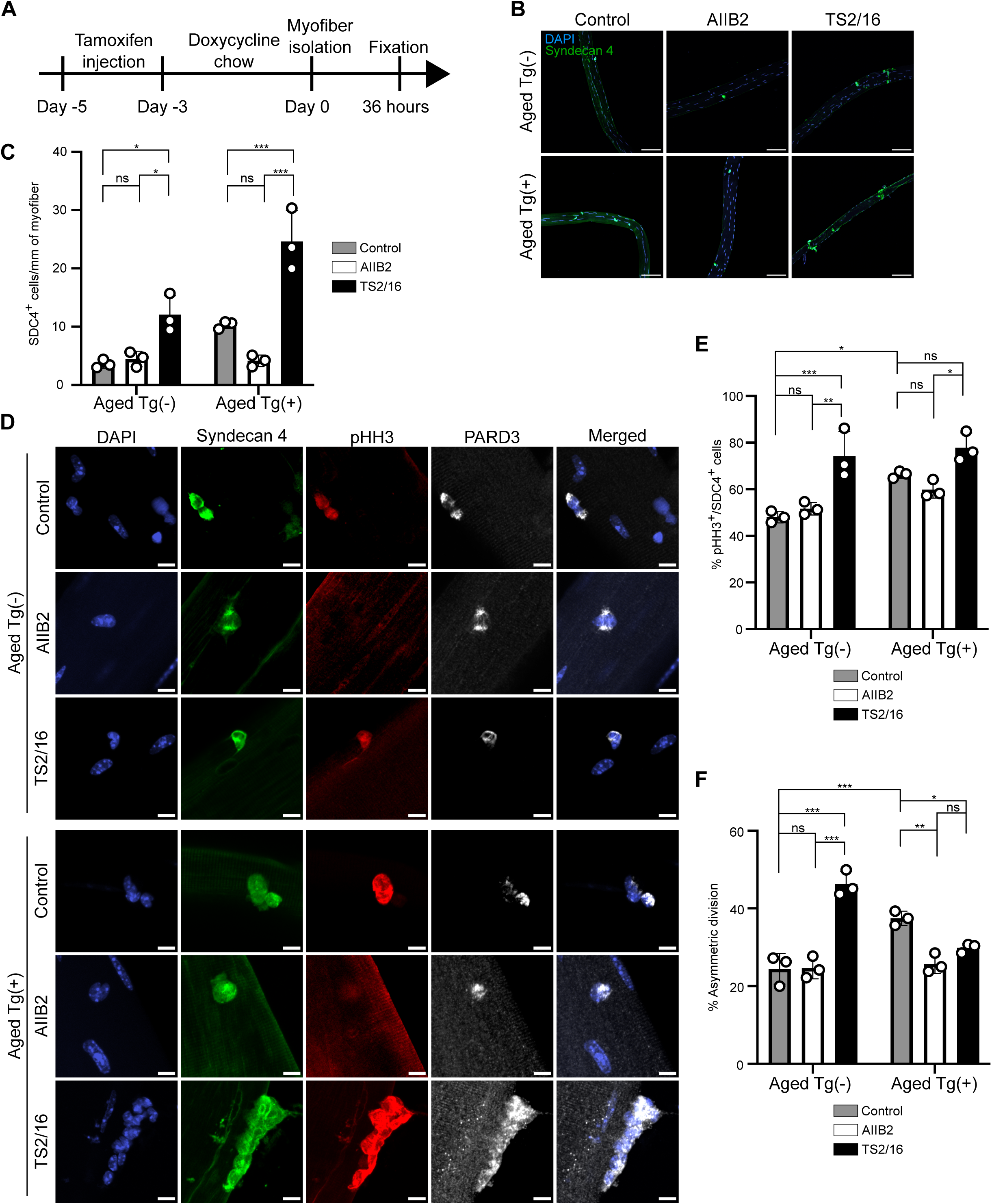
Activating FGFR1 restores SC sensitivity of Integrin-β1 to regulate asymmetric division in SCs from aged mice. (A) Experimental scheme showing treatment of aged caFGFR1– and caFGFR1+ mice and the timeline for myofiber harvest and culture. (B) Representative images of myofibers from aged caFGFR1–and caFGFR1+ mice cultured in the presence of AIIB2 antibody or TS2/16 antibody, where SCs were identified by Syndecan-4 immunoreactivity. Scale bars, 100 μm. (C) Syndecan-4^+^ cells on myofibers quantified from aged caFGFR1– and caFGFR1+ mice cultured in the presence of AIIB2 or TS2/16. *n* > 10 myofibers analyzed with samples from 3 mice, and statistics performed based on 3 independent mice; data are presented as mean ± SD; *\*P* < 0.05 and *\*\*\*P* < 0.001 in a two-way ANOVA test. (D) Representative images of SCs on myofibers from aged caFGFR1– and caFGFR1+ mice cultured in the presence of AIIB2 antibody or TS2/16 antibody, where SCs were identified by Syndecan-4 immunoreactivity and assayed for pHH3 immunoreactivity and PARD3 immunoreactivity. Scale bars, 10 μm. (E and F) The percentage of pHH3^+^ cells (E) and percentage of asymmetric division (F) on myofibers from aged caFGFR1– and caFGFR1+ mice cultured in the presence of AIIB2 or TS2/16. *n* > 200 Syndecan-4^+^ cells analyzed with samples from 3 mice, and statistics performed based on 3 independent mice; data are presented as mean ± SD; *\*P* < 0.05, *\*\*P* < 0.01, and *\*\*\*P* < 0.001 in a two-way ANOVA test.

Syndecan-4^+^ SCs were assessed for pHH3 and PARD3 immunoreactivity (**Figure 6D**). When quantified, 48 ± 2% of the Syndecan-4^+^ cells on myofibers from aged caFGFR1– mice were pHH3^+^, with no significant change occurring following AIIB2 treatment (**Figure 6E**). Syndecan-4^+^ cells on myofibers from aged caFGFR1–mice increased to 74 ± 11% following TS2/16 treatment. On myofibers from aged caFGFR1+ mice, the percentage of pHH3^+^/Syndecan-4^+^ cells increased to 66 ± 2%, but was unchanged following AIIB2 treatment, and rose to 78 ± 6% following TS2/16 treatment.

Activating FGFR1 signaling in aged mice drives asymmetric cell division^6^, and thus, we queried whether activating or inhibiting Integrin-β1 affects the numbers of SCs dividing asymmetrically. On myofibers from aged caFGFR1– mice, 25 ± 4% of dividing were asymmetric, with no change in asymmetrically dividing cells following AIIB2 treatment (**Figure 6F** and **Figure S4B**). However, the number of asymmetrically dividing Syndecan-4^+^ cells increased to 46 ± 3% following TS2/16 treatment, restoring levels comparable SCs from young mice suggesting that activating Integrin-β1 without exogenous FGF promotes asymmetric division in SCs from aged mice. If FGFR1 signaling was activated in aged caFGFR1+ mice, asymmetric division of SCs increased to 38 ± 2%, similar to levels in young mice. However, AIIB2 treatment decreased asymmetrically dividing SCs to 26 ± 2%, demonstrating involvement of Integrin-β1 in regulating SC asymmetric division. Unexpectedly and in contrast to caFGFR1– mice, TS2/16 treatment of myofibers from aged caFGFR1+ mice did not increase SC asymmetric division above the levels occurring in caFGFR1+ mice, demonstrating that activating Integrin-β1 when FGFR1 is ectopically activated promotes SC symmetric division. Thus, Integrin-β1 and FGFR1 coordinate to regulate SC asymmetric division or symmetric division and appear to affect signaling pathways that are compromised in aged mice.

### Constitutive activation of FGFR1 in SCs from aged mice restores their responsiveness to RGD density on myofibers in hydrogels

FGFR1 and Integrin-β1 appear to cooperatively regulate SC asymmetric division and symmetric expansion. Therefore, we asked if ectopically activating FGFR1 would restore the age-dependent loss of SC sensitivity to RDG density. To determine whether constitutive FGFR1 activation rescues SC responsiveness to RGD density on myofibers from aged mice, myofibers isolated from aged caFGFR1– and caFGFR1+ mice were encapsulated in viscoelastic hydrogels containing either 0.1 mM or 1 mM RGD **(Figure 7A)**. Mice received daily tamoxifen injections for two consecutive days, followed by three days of doxycycline chow prior to myofiber isolation. The following day, isolated myofibers were encapsulated and cultured for 2 days, with a 2-hour EdU pulse administered before harvest to assess SC proliferation. SC fate was evaluated by Pax7 and MyoD immunoreactivity **(Figures 7B)**.

**Figure 7.**
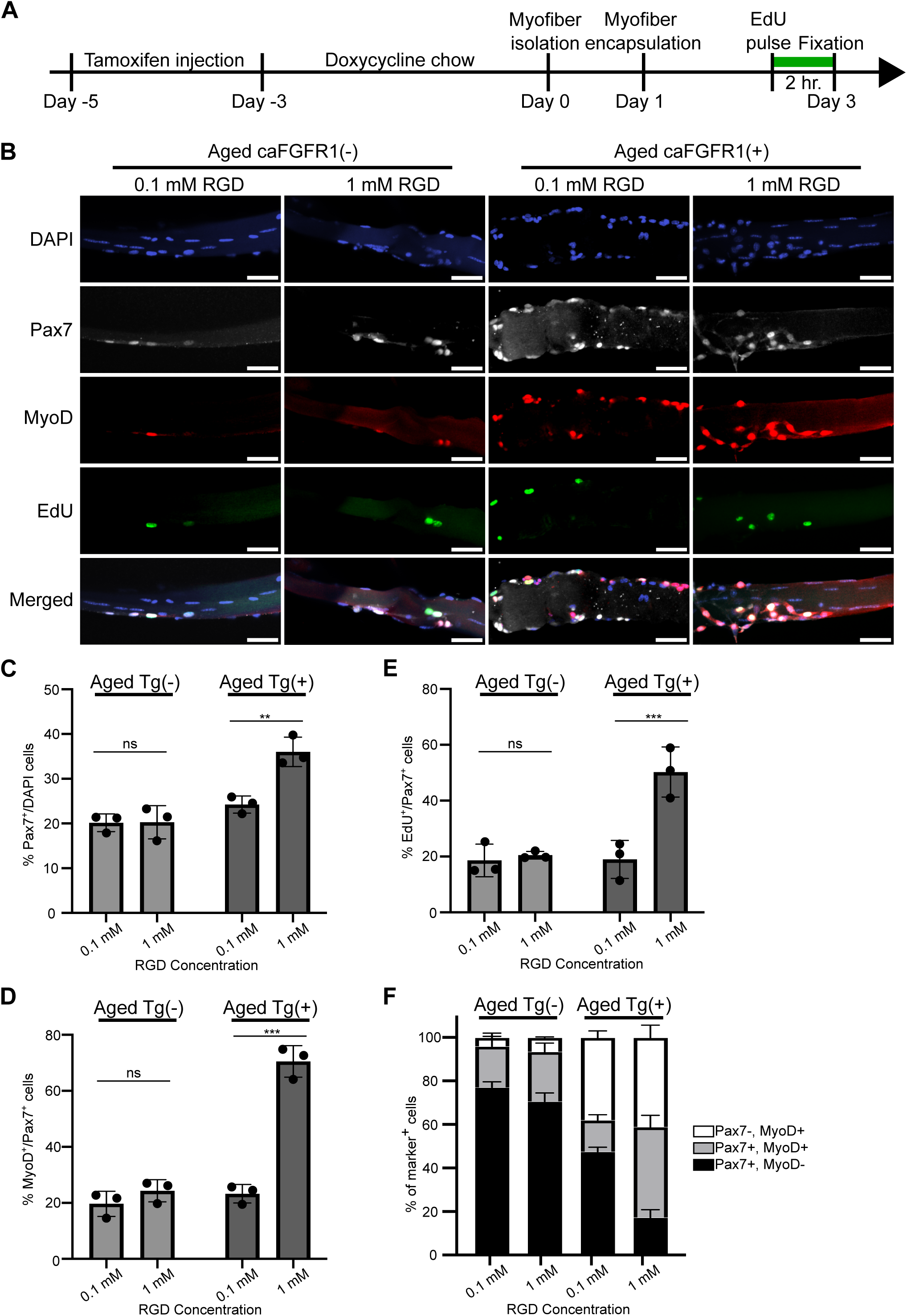
Constitutively active FGFR1 in SCs on myofibers from aged mice restores responsiveness to RGD density in hydrogel cultures. (A) Experimental scheme showing treatment of aged caFGFR1– and caFGFR1+ mice. (B) Representative images of myofibers from aged caFGFR1– and caFGFR1+ mice embedded in hydrogels with 0.1 mM and 1 mM RGD and assessed for Pax7, EdU, and MyoD immunoreactivity after 2 days of culture. Scale bars, 50 μm. (C-E) Quantification of Pax7^+^ (C), MyoD^+^ (D), and EdU^+^ (E) for SCs on myofibers from aged caFGFR1– and caFGFR1+ mice embedded in hydrogels with 0.1 mM and 1 mM RGD. *n* > 9 myofibers analyzed with samples from 3 mice, and statistics performed based on 3 independent mice; data are presented as mean ± SD; *\*\*P* < 0.01 and *\*\*\*P* < 0.001 in a two-way ANOVA test. (F) Percentage of Pax7+/MyoD–, Pax7+/MyoD+, and Pax7–/MyoD+ cells on myofibers from aged caFGFR1– and caFGFR1+ mice embedded in hydrogels with 0.1 mM and 1 mM RGD. *n* > 9 myofibers analyzed with samples from 3 mice, and data are presented as mean ± SD.

Fewer than 25% of cells on myofibers from aged caFGFR1– mice were Pax7^+^, regardless of RGD concentration **(Figures 7C)**. In contrast, on myofibers from aged caFGFR1+ mice in 0.1 mM RGD hydrogels, 24 ± 2% of cells were Pax7^+^, increasing to 36 ± 3% Pax7^+^ cells in 1 mM RGD. Thus, high RGD density supports SC maintenance in aged mice when FGFR1 is ectopically activated. Among Pax7^+^ cells from aged caFGFR1– mice, fewer than 25% expressed MyoD regardless of RGD concentration **(Figures 7D)**. On myofibers from aged caFGFR1+ mice, MyoD^+^/Pax7^+^ cells increased from 23 ± 3% in hydrogels with 0.1 mM RGD to 71 ± 6% in hydrogels with 1 mM RGD. Fewer than 25% of Pax7^+^ cells were EdU^+^ on myofibers from aged caFGFR1– mice with no response to RGD concentration changes **(Figure 7E)**. However, EdU incorporation in SCs from aged caFGFR1+ mice increased from 19 ± 7% in hydrogels with 0.1 mM RGD to 50 ± 9% in 1 mM RGD.

On myofibers from aged caFGFR1– mice, over 70% of SCs remained quiescent (Pax7+/MyoD–), ∼20% were activated SCs (Pax7+/MyoD+), and few were differentiating myoblasts (Pax7–/MyoD+), with no significant differences between high and low RGD densities **(Figure 7F)**. In contrast, on myofibers from aged caFGFR1+ mice in 0.1 mM RGD hydrogels, the number of quiescent SCs was 48 ± 2%, with 14 ± 2% activated SCs and 38 ± 3% differentiating myoblasts. When encapsulated in 1 mM RGD hydrogels, the numbers of quiescent SCs further declined to 17 ± 4%, while activated SCs increased to ∼40% and differentiating myoblasts increased to ∼40%. Activating FGFR1 in SCs from aged mice appears to restore their responsiveness to RGD density, and at high RGD density in the presence of ectopically activated FGFR1, SCs exit from quiescence and expand.

## Discussion

SCs are indispensable for muscle regeneration, and thus, their functional decline with age contributes to sarcopenia and impaired recovery from injury^9, 25, 26^. The decline in SC function is partially attributed to the reduced abundance of fibronectin, a key extracellular matrix protein that supports SC maintenance and regeneration of muscle^18^. We employed a fully defined synthetic viscoelastic hydrogel with tunable densities of fibronectin-derived RGD ligands to encapsulate SCs with their associated myofibers to assess how SCs from aged mice respond to fibronectin in a controlled environment^19, 27^. The synthetic hydrogel or ECM platform mimics muscle-relevant viscoelastic microenvironments while preserving SC-myofiber interactions occurring *in vivo* that maintains quiescent (Pax7+/MyoD-) SCs and SC polarity. SCs from aged mice embedded in the hydrogel ECM platform respond poorly to variations in RGD density compared to SCs from young mice, identifying an age-related decline in sensitivity of SCs to fibronectin-mediated cues that may contribute to loss of SC self-renewal and declines in SC function occurring during aging.

Sensing of fibronectin by SCs correlates with co-localization of Integrin-β1, Syndecan-4, and FGFR1, suggesting that all three membrane proteins may be involved. Integrin-β1 is the primary receptor mediating SC adhesion to fibronectin and is critical for maintaining self-renewal capacity^11^, while Syndecan-4 is required for FGF binding and signaling via FGFR1^13, 14^. The loss of fibronectin sensitivity does not arise from lower levels of protein as we observed no significant differences in Syndecan-4 or Integrin-β1 protein abundance during aging. During aging, FGFR1 signaling is compromised, reducing SC self-renewal, which is recovered by ectopically activating FGFR1 in the absence of FGF^6^ (Cutler et al., unpublished). Syndecan-4 is required for FGF-2 binding to FGFR1^14^ and Integrin-β1 enhances signaling^11^, both of which are compromised in SCs from aged mice. We hypothesized that impaired fibronectin sensing in SCs from aged mice arises from a failure of FGFR1 to functionally coordinate with Integrin-β1 and Syndecan-4. SCs from aged mice generally fail to polarize FGFR1^6^, and we demonstrate this is accompanied by a failure to co-localize Integrin-β1 and Syndecan-4 with FGFR1. Ectopically activating FGFR1 rescues co-localization of Syndecan-4 and Integrin-β1 with FGFR1 as well as asymmetric division, positioning FGFR1 as a critical receptor coordinating matrix receptor activity via integrating structural adhesion and growth factor signaling. This coordination is especially critical during regeneration, when extracellular fibronectin is upregulated and functions as a key instructive niche component.

FGF signaling, essential for SC self-renewal^8^, requires functional Integrin-β1as disrupting either Integrin-β1 or FGFR1 impairs SC quiescence and the capacity of SCs to repair muscle^6, 11^. Activating Integrin-β1 in the absence of exogenous FGF is sufficient to promote SC asymmetric division and preserve the SC pool, underscoring the role of adhesion-mediated cues in SC maintenance. However, simultaneous activation of Integrin-β1 and FGFR1 shifts dividing SCs to symmetrically expand, suggesting that the cooperative signals regulate whether the SC pool is maintained or amplified. The degree of SC self-renewal or SC expansion may temporally regulate SCs numbers as the muscle regenerates following an injury to meet regenerative demand. During aging, impaired FGFR1 signaling in SCs is similar to SCs from young mice in the absence of added FGF as SCs fail to maintain asymmetric division^6^. Constitutively activating FGFR1 in SCs from aged mice restores the capacity for asymmetric expansion and responsiveness to RGD density. Either activating Integrin-β1 or exposing SCs from aged mice to a high-RGD environment when FGFR1 is constitutively active promotes SC to symmetrically expand. These findings confirm and extend prior reports independently proposing that aging impairs extrinsic niche cues and intrinsic signaling pathways governing SC behavior^9, 26^. By integrating stem cell niche cues arising from Integrin-β1 with FGFR1-mediated signaling, we show that RGD concentrations and the levels of Integrin-β1 signaling critically regulate SC stem cell numbers, balancing self-renewal and expansion. Therefore, we envision that biomaterials co-presenting integrin ligands and transiently sustained FGFR signals promote SC expansion *ex vivo* and have the capability to improve regenerative potential when implanting hydrogels, embedded SCs, or skeletal muscle organoids into aged muscle.

### Limitations of study

Our study focused on the co-dependence of fibronectin and FGFR1 signaling, and thus, it is possible that other ECM ligands, including collagen and laminin, are remodeled as muscle regenerates and may be similarly compromised during aging. Moreover, FGFR4 is expressed in SCs and involved in maintaining quiescence^28^ but was not studied in this series of experiments. The impact of ECM compositional changes on SCs in aged mice is largely unexplored. We have developed a platform whereby peptides derived from these proteins could be systematically incorporated into hydrogel microenvironments to assess their singular or combinatorial effects on SC behavior in aged mice. Although we identify a causal role for Integrin-β1 and FGFR1 in modulating asymmetric division, the downstream pathways remain incompletely defined as signaling pathways involving p38α/β MAPK^8^, the TTP family of RNA binding proteins^29^, AUF1^30^, microRNAs^31, 32^, Wnt7a^33^, and EGF^34^ signaling appear involved. Further studies are needed to dissect the roles of these signals and the mechanisms that integrate signaling decisions with symmetric expansion or asymmetric division of SCs. While encapsulation of myofibers in hydrogels, coupled with real-time tracking of SCs, could serve as an *in vitro* model to study SC behavior, *in vivo* studies are needed to determine whether biomaterials presenting integrin ligands, in combination with pharmacologic restoration of FGFR1 activity, can rejuvenate SC function in aged mice and improve muscle maintenance and repair in aged organisms.

## Resource availability

### Lead contact

Further requests for resources and information should be directed to and will be fulfilled by the lead contact, Bradley B. Olwin.

### Materials availability

All materials generated in this study are available from the lead contact with a completed materials transfer agreement.

### Data and code availability

All data reported in this paper will be shared by the lead contact upon request. This paper does not report original code. Any additional information required to reanalyze the data reported in this paper is available from the lead contact upon request

## Acknowledgments

This work was supported by grants from the NIH (DE016523 and DK120921) to KSA and NIH (AR049446 and AR070630) to BBO. The imaging work was performed at the BioFrontiers Institute’s Advanced Light Microscopy Core (RRID: SCR_018302). Laser scanning confocal microscopy was performed on a Nikon A1R microscope supported by NIST-CU Cooperative Agreement award number 70NANB15H226 and a Nikon AXR microscope supported by NIH Grant 1S10OD034320. The data analysis and visualization work was conducted using the Imaris Analysis Workstation supported by NIH 1S10RR026680-01A1.

## Author contributions

Conceptualization, T.L.C., B.B.O., and K.S.A.; Methodology: T.L.C., A.A.C., and T.K.V.; Software: T.S.Z., A.A.C., and J.V.K.; Formal analysis: T.L.C., T.S.Z., and T.K.V.; Investigation: T.L.C., T.S.Z., C.H.B, and T.K.V.; Supervision: B.B.O. and K.S.A.; Writing—original draft: T.L.C. and T.S.Z.; Writing—review & editing: T.L.C., T.S.Z., T.K.V., B.B.O., and K.S.A.

## Supplemental information

Document S1. Figures S1-S4 and Table S1-S2

## STAR Methods

### Animals

All mice were bred and housed according to National Institutes of Health (NIH) guidelines for the ethical treatment of animals in a pathogen-free facility at the University of Colorado at Boulder. All animal protocols and procedures were approved by the University of Colorado Institutional Animal Care and Use Committee, and the conducted studies complied with all ethical regulations. Wild-type mice were C57BL/6 (Jackson Labs, ME, USA). Pax7^iresCre^; ROSA26^LSLrtTA^; caFGFR1 mice were generated by crossing C57BL/6, Pax7^iresCre^ mice^35^, ROSA26-rtTA mice (Jackson Labs)^36^, and conditional caFGFR1 transgenic mice^22^. Mice from 4-6 months of age were considered young, and mice from 20-24 months of age were considered aged. Experiments were restricted to male mice only. Genotyping was performed on tissue samples collected at weaning and sent to Transnetyx for automated genotyping.

### Animal procedures

Mice were anesthetized with isoflurane, followed by injection of 50 μL 1.2% BaCl₂ into the left tibialis anterior (TA) muscle, and then myofibers from both injured and contralateral uninjured muscles were harvested 7 days post-injury. Recombination was induced by intraperitoneal injection of tamoxifen (2 mg/mouse in corn oil; Sigma-Aldrich) for 2 consecutive days, followed by doxycycline chow (Envigo TD.01306) for 3 days prior to myofiber isolation

### Software Packages

- CellRanger Software Suite/3.0.1
- FastQC 0.11.8
- R 4.3.3
- Seurat 5.1.0
- CellChat 1.6.1
- SoupX 1.5.2

### Quality control, read alignment, and expression quantification

*FastQC* was used to evaluate the quality of read depth on each replicate of Fastq files from sequencing. *CellRanger* was employed to process Fastq files by aggregating technical replicates and generating gene count matrices. Transcriptomes from snRNA-seq were aligned using a previously validated custom pre-mRNA mm10 reference package (Cutler et al., unpublished).

### Accounting for experimental noise and doublets

*CellRanger*-aligned feature matrices were loaded into *R* using the load10X() function from the *SoupX* package. AutoEstCont() was used to predict RNA contamination and adjustcounts() normalized counts to an estimated noise parameter. Manual correction of gene expression matrices removed doublets and debris using recommended metrics. In addition, cells or nuclei expressing more than 2,500 or less than 200 unique features were removed.

### Normalization, dimensional reduction, and nuclear clustering

The tutorial provided by the Mayaan lab (satijalab.org/seurat/articles/pbmc3k_tutorial) was followed to normalize data and mitochondria and low-quality nuclei were removed. Seurat objects were processed using the NormalizeData(), FindVariableFeatures(), and ScaleData() functions from *Seurat* to scale, and log normalize gene counts within each sample. Then, the *Seurat* functions FindIntegrationAnchors() and IntegrateData() were used to integrate data using reciprocal PCA (rPCA). Nuclear clusters were separated using the shared nearest neighbor (SNN) modularity optimization-based clustering. The minimum number of neighbors, minimum distance between neighbors, and resolution parameters were adjusted while performing dimensional reduction of the integrated *Seurat* object using UMAP.

### CellChat Analysis of Cell-Cell Communication

Cell-cell communication analysis of each *Seurat* object was performed using *CellChat* (v1.6.1) following the workflow^37^. ComputeCommunProb() was used with an 18% Truncated Mean cutoff to determine the cell-type specific expression of ligand or receptors in a pathway of interest.

### Myofiber isolation

Extensor digitorum longus muscles were dissected and digested in Ham’s F-12 (Gibco) containing collagenase (400 U/mL) (Worthington) at 37 °C for 1.5 hours with gentle rotation. After digestion, the collagenase was inactivated with Ham’s F-12 containing 15% horse serum (Gibco) and 0.8 mM CaCl_2_. Then, a flame-polished glass pipet was used to transfer individual myofibers to 6-well tissue culture plates containing Ham’s F-12 supplemented with 15% horse serum, 1% penicillin-streptomycin (Gibco), and 0.8 mM CaCl_2_. For genetically engineered mice, medium was supplemented with 1 µg/mL doxycycline. When indicated, myofibers were either fixed in 4% paraformaldehyde for 10 min immediately after isolation or cultured in suspension at 37 °C and 5% CO₂ for 36 h prior to fixation or 24 h prior to encapsulation, with supplementation of 0.5 nM fibroblast growth factor 2, 1 µg/mL AIIB2 (Developmental Studies Hybridoma Bank), or 1 µg/mL TS2/16 (eBioscience) as specified.

### Immunocytochemistry of myofibers

Fixed myofibers were permeabilized with 0.25% Triton-X100 and 2% bovine serum albumin (BSA) in phosphate buffered saline (PBS, pH 7.4) for 60 min at room temperature (RT). For photo-expanded samples, myofibers were blocked with 2% BSA in PBS for 60 min at RT. Samples were washed three times for 5 min each in PBS and incubated overnight at 4 °C with primary antibodies diluted in PBS containing 2% BSA. After three 5 min washes with PBS, myofibers were incubated with Alexa secondary antibodies (Thermo Fisher Scientific, 1:500) and 4′,6-diamidino-2-phenylindole (DAPI, 1 μg/mL) in PBS containing 2% BSA for 1 hour at RT. Primary antibodies included chicken anti-Syndecan-4^10^ at 1:200, rat anti-mouse CD29 (BD Biosciences, 9EG7) at 1:200, rabbit anti-FGFR1 (abcam, ab10646) at 1:200, rabbit anti-FGFR3 (Sigma-Aldrich, F3922), rat anti-phospho-H3 (Sigma-Aldrich, H9908) at 1:200, and rabbit anti-PARD3 (Bio-Techne, NBP1-88861). Myofibers were transferred onto glass slides, mounted in Mowiol supplemented with DABCO (Sigma-Aldrich), and covered with coverslips.

### Synthesis of HA-hydrazide and characterization

HA-hydrazide was synthesized as previously described^27, 38^. Briefly, HA (60 kDa, 500 mg) was dissolved in dH₂O (100 mL), followed by the addition of adipic acid dihydrazide (≈6.5 mg, >60-fold molar excess) while adjusting the pH to 6.8. 1-Ethyl-3-(3-dimethylaminopropyl)carbodiimide (EDC, 776 mg) and hydroxybenzotriazole (HOBt, 765 mg), separately dissolved in a 1:1 DMSO/dH₂O mixture, were added dropwise to the reaction. The pH was maintained at 6.8 every 30 min for 4 h, and the reaction proceeded for 24 h. The product was dialyzed against dH₂O (8000 MWCO) for 3 days and lyophilized for 3 days. Then the product was redissolved in 5 wt.% NaCl/dH₂O, precipitated in ethanol, redialyzed for three days, and lyophilized for an additional three days. The final white powder (430 mg, 86% yield) was flash frozen in liquid nitrogen and stored at −20 °C. Molecular weight (∼86 kDa) was confirmed by GPC characterization and functionalization (∼28.5%) was confirmed by ¹H NMR^19^.

### Synthesis of HA-aldehyde and characterization

HA-aldehyde was synthesized as previously described^27, 38^. Briefly, HA (500 kDa, 500 mg) was dissolved in dH_2_O (50 mL), and then sodium periodate (267.5 mg) was added into the reaction mixture. After stirring in the dark for 2 h, the reaction was quenched with ethylene glycol (70 µL), dialyzed against dH₂O (8000 MWCO) for 3 days, and lyophilized for another 3 days to obtain a white powder (471 mg, 94% yield). HA-aldehyde was flash frozen in liquid nitrogen and stored at −20 °C. The functionalization of the HA-aldehyde macromer was quantified using a 2,4,6-trinitrobenzene sulfonic acid (TNBS) assay as previously describe^39, 40^. Briefly, the HA-aldehyde was dissolved at 2 wt.% (w/v) in dH₂O and reacted with tert-butyl carbazate (t-BC, in 1% trichloroacetic acid). After 24 h, the HA-aldehyde/t-BC and t-BC standards were reacted with 0.5 mL of TNBS solution (6 mM in 0.1 M sodium tetraborate, pH 8) for 1 h. The reaction was then quenched with 0.5 N hydrochloric acid. Molecular weight (∼148 kDa) was confirmed by GPC characterization and functionalization (∼36%) was confirmed by absorbance measured at 340 nm using a microplate reader^19^.

### PEG-bicyclononyne synthesis

PEG-bicyclononyne (BCN) was synthesized as previously described^27, 41^. Briefly, 8-arm PEG-amine (40 kDa, 1.0 g, 0.2 mmol amine) and BCN-oSu (0.1 g, 0.343 mmol) were added to a 50 mL round-bottom flask. The components were dissolved in anhydrous DMF (10 mL), and the solution was stirred at room temperature under argon. N,N-Diisopropylethyleamine (0.8 mmol) was added, and the reaction proceeded overnight. The mixture was then diluted with dH_2_O and dialyzed for 3 days (8000 MWCO). The product was lyophilized for 3 days to yield a white powder (0.985 g, 98% yield). End-group functionalization (>95%) was confirmed by ¹H NMR^19^.

### Hydrogel Formulation

Hydrogels were prepared by dissolving functionalized HAs in PBS at 5 wt.%. Functionalized PEG-BCN was dissolved in PBS at 10 wt.%, and the azide-PEG_3_-phenolaldehyde (BroadPharm) crosslinker was prepared at 20 mM in PBS. Final hydrogels were formed with a polymer content of 5 wt.%, based on stoichiometric calculations. In the reported formulation, 12% of the hydrazide groups on HA were substituted with azide–PEG₃–phenolaldehyde, yielding a hydrogel composed of 88% adaptable alkyl-hydrazone crosslinks and 12% irreversible azide-alkyne crosslinks, with PEG-BCN and HA-aldehyde incorporated at corresponding molar ratios.

### Myofiber encapsulation

The hydrogels were formed by pre-reacting the 5wt.% HA-hydrazide with the azide-PEG_3_-phenolaldehyde (20 mM) and the benzaldehyde-KRGDS in PBS overnight at 4°C. The HA-aldehyde, PEG-BCN and PBS solutions were mixed. Myofibers were picked and placed into a glass-bottom 96-well plate (Cellvis) using a flame-polished glass pipet (approximately 15 myofiber per well). After allowing the myofibers to settle for 10 minutes in an incubator, excess medium was carefully removed. Immediately after media removal from the myofibers, the HA-hydrazide mixture was added, and then the HA-aldehyde mixture was mixed into the myofibers with HA-hydrazide solution. Myofibers were gently resuspended in 30 μL of hydrogel solution. After polymerization for 20 minutes in the incubator, fresh Ham’s F-12 supplemented with 15% horse serum, 1% penicillin-streptomycin, and 0.8 mM CaCl_2_ was added. For genetically engineered mice, medium was supplemented with 1 µg/mL doxycycline. Supplementation of 0.5 nM fibroblast growth factor 2 was added as indicated. Encapsulated myofibers were cultured at 5% CO_2_ and 37 °C in the incubator.

### Myofiber proliferation assays and immunocytochemistry

Hydrogels containing myofibers were treated with 10 μM EdU (Thermo Fisher Scientific) for 2 hours after two days of culture. Then, the samples were quickly washed with PBS and fixed with 10% formalin for 30 minutes. After three 10 min PBS washes, the samples were permeabilized with 0.5% Triton X-100 and 4.6 mM N_3_-PEG_3_-OH in PBS for two hours. The samples were washed with PBS, and then incubated with the Click-iT reaction cocktail using the Click-iT EdU Alexa Fluor 488 kit (Thermo Fisher Scientific) for 30 minutes. After three PBS washes, the samples were incubated with 20 mM NH_4_Cl in PBS for 1 hour. All samples were then washed with PBS and blocked with 5% BSA in PBS at 4 °C overnight. The samples were immunolabeled with Pax7 (mouse, Developmental Studies Hybridoma Bank, 1:250) and MyoD (rabbit, Santa Cruz Biotechnology, SC-760, 1:250) primary antibodies in PBS containing 5% BSA at 4 °C overnight. After washing with PBS, samples were incubated with Alexa secondary antibodies (Thermo Fisher Scientific, 1:500) and DAPI (1 μg/mL) in PBS containing 5% BSA at 4 °C overnight. All samples were washed three times with PBS for 10 minutes prior to imaging.

### Photo-expansion microscopy

Following incubation with secondary antibodies, myofibers were washed three times with PBS and treated with 0.1 mg/ mL acryloyl-X in PBS at 4 °C overnight. After PBS washes, myofibers were transferred onto a glass slide using a flame-polished glass pipet, and excess PBS was carefully removed. 10 μL of the PhotoExM formulation **(Table S2)** was applied to cover the myofibers and a Rain-X-coated coverslip was gently placed on top, spaced 400 μm apart using rubber gaskets. Samples were incubated in the dark at room temperature for 15 minutes. The samples were then polymerized with *λ* = 365 nm, *I* = 4.5 mW/cm^2^ light for 70 seconds, followed by overnight digestion at 40 °C in PBS containing Proteinase K (8 U/mL). After digestion, the samples were expanded in dH_2_O for 20 minutes. The samples were then stained with DAPI in dH_2_O for 1 hour and expanded two more times in dH_2_O for 20 minutes each. Before imaging, the hydrogels were mounted on glass bottom dishes with poly-L-lysine.

### Imaging and image analysis

Images of standard immunostained samples were carried out using a Nikon A1R Confocal Microscope equipped with a x20 NA = 0.75 objective. Images were visualized and analyzed with ImageJ. Images of expanded samples were carried out using a Nikon AXR Confocal Microscope equipped with a x60 NA = 1.2 water immersion objective. Images were visualized and analyzed with Imaris. The brightness and contrast of each channel was separately adjusted for better visualization.

### Statistical analysis

Statistics were analyzed using Prism (GraphPad), and *P* < 0.05 was considered significant. For hydrogel-encapsulated myofibers, at least 9 myofibers were analyzed from 3 independent mice, and two-way ANOVA tests were performed based on the three biological replicates. For transcript levels in SCs from single-cell and single-nucleus sequencing data, statistical significance was measured by *Seurat’s* Wilcox rank sum test. For protein expression levels in SCs on myofibers, at least 10 myofibers were analyzed from 3 independent mice, and one-way ANOVA tests were performed. For percentage of SDC4^+^ cells with polarized FGFR1, at least 60 cells were analyzed from 3 independent mice, and a one-way ANOVA test was performed based on the three biological replicates. For FGFR polarization and co-localization analysis, at least 10 cells were analyzed from 3 independent mice, and one-way ANOVA tests were performed based on the three biological replicates. For quantification of SDC4^+^ cells, pHH3^+^ cells, and asymmetric division, at least 200 cells from a minimum of 10 myofibers were analyzed from 3 independent mice, and two-way ANOVA tests were performed based on the three biological replicates.

### Graphical illustration

Schematic illustrations of hydrogels and photo-expansion microscopy were created with BioRender.com.

