## Supplemental for "Activating FGFR1 restores Integrin-β1–mediated fibronectin sensing in satellite cells of aged mice"

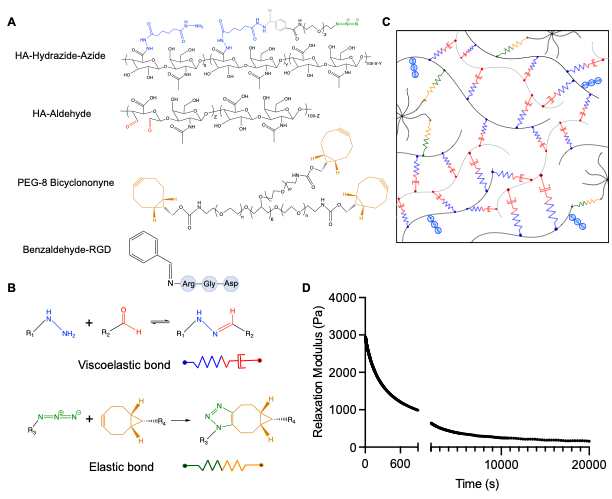

**Figure S1. Network structure and rheological properties of the hydrogel, related to Figure 1.** (A and B) Chemical structures (A) of 86 kDa HA-hydrazide-azide, 148 kDa HA-aldehyde, and 40 kDa 8-arm PEG-BCN, and schematic representations of the functional groups (B) forming network crosslinks in the hydrogel, including alkyl-hydrazone (viscoelastic) and triazole (elastic) bonds. (C) Schematic representation of the HA-PEG hybrid hydrogel composed of 88% adaptable alkyl-hydrazone crosslinks and 12% irreversible azide-alkyne crosslinks, forming a stable, stress relaxing network. (D) Stress relaxation profile of the hydrogel, showing stress relaxation from an initial modulus of ~3000 Pa to ~1200 Pa at 600 seconds, ultimately reaching a final modulus of ~180 Pa.

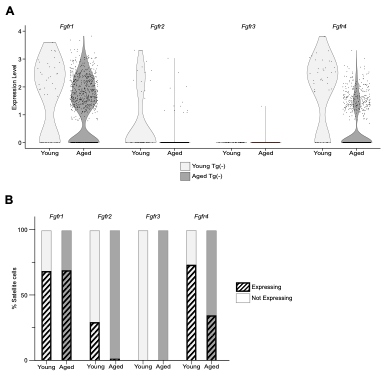

**Figure S2. Satellite cells from young and aged caFGFR1– mice do not express *Fgfr3* transcripts, related to Figure 4.** (A) Violin plots showing transcript expression levels of *Fgfr1*, *Fgfr2*, *Fgfr3*, and *Fgfr4* in SCs from young and aged caFGFR1– mice, derived from single-cell and single-nucleus RNA sequencing datasets. (B) Percentage of satellite cells from young and aged caFGFR1– mice expressing *Fgfr1*, *Fgfr2*, *Fgfr3*, or *Fgfr4* transcripts.

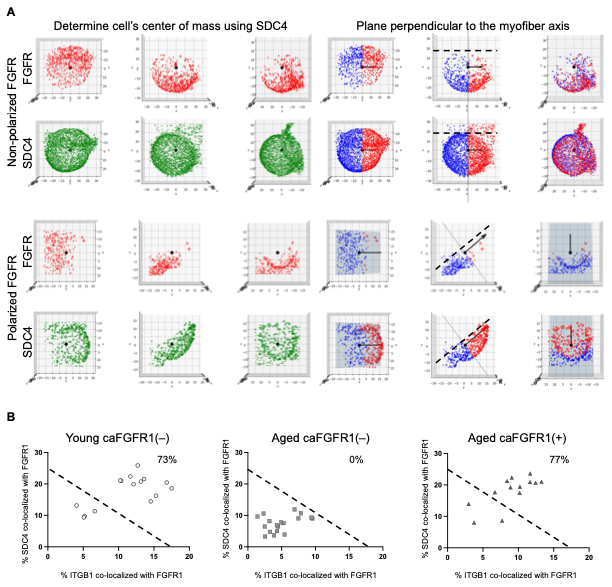

**Figure S3. FGFR1 polarization and co-localization with syndecan-4 and integrin-β1 are reduced in SCs from aged mice and restored by constitutive FGFR1 activation, related to Figure 4.** (A) Representative images of SDC4^+^ SCs exhibiting non-polarized or polarized FGFR1, illustrating the method used to quantify FGFR1 polarization. The center of mass of each SC was determined from the SDC4 signal, and a plane perpendicular to the myofiber axis (dashed line) was drawn through this center. FGFR1 volume on each side of the plane was summed and expressed as a ratio of the total FGFR1 volume within the cell. These ratios are represented as (0.5 ± *x*), where *x* ranges from 0 to 0.5. The variable *x*, indicating the fraction of FGFR1 redistributed across the plane, was normalized to a scale from 0 to 1 to define the degree of FGFR1 polarization. (B) Correlation plots showing co-localization of SDC4 (y-axis) or ITGB1 (x-axis) with FGFR1 in SCs on myofibers from young caFGFR1–, aged caFGFR1–, and aged caFGFR1+ mice. Each point represents a single SC measurement. The upper-right quadrant indicates SCs with relatively high levels of both ITGB1 and SDC4 co-localized with FGFR1 with the percentage of cells within this quadrant quantified and indicated.

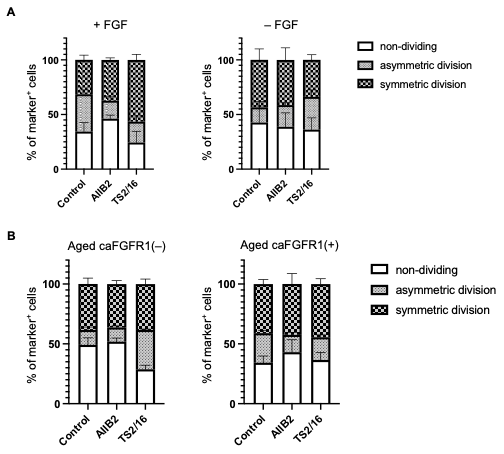

**Figure S4. Percentage of SDC4^+^ cells undergoing symmetric division, asymmetric division, or not dividing, related to Figure 5 and 6.** (A) Percentage of SDC4^+^ cells undergoing symmetric division, asymmetric division, or not dividing on myofibers from young caFGFR1– mice, cultured with or without FGF-2, in the presence of AIIB2 or TS2/16. (B) Percentage of SDC4^+^ cells undergoing symmetric division, asymmetric division, or not dividing on myofibers from aged caFGFR1– and caFGFR1+ mice, cultured in the presence of AIIB2 or TS2/16.

**Table S1. Mechanical properties of the hydrogel, related to Figure 1.**

| **Property** | **Value** |
| --- | --- |
| Shear storage modulus after formation (Pa) | 2313 ± 560 |
| Equilibrium swollen shear storage modulus (Pa) | 803 ± 114 |
| Stress relaxation time constant (min) | 56 ± 3 |
| Maximum stress relaxed (%) | 94 ± 3 |
| Stress relaxed after 10 min (%) | 57 ± 6 |

*Note.* It has been reported that a mouse gastrocnemius muscle relaxes roughly 70% of the applied stress within 600 seconds.

**Table S2. PhotoExM formulations used for imaging myofibers with ~4.5X linear expansion, related to Figure 4.**

| **Full name** | **Supplier** | **Concentration** |
| --- | --- | --- |
| Sodium acrylate | Combi-Blocks | 16 wt.% |
| Poly(ethylene glycol)-diacrylamide  MW 600 g/mol | Creative PEGWorks | 0.875 wt.% |
| 8-arm, Poly(ethylene glycol), thiol functionalized  MW 10000 g/mol | Jenkem | 6 wt.% |
| Acrylamide | Sigma Aldrich | 3 wt.% |
| Lithium phenyl-2,4,6-trimethylbenzoylphosphinate | Sigma Aldrich | 0.2 wt.% |
